# Sperm production and allocation in response to risk of sperm competition in the black soldier fly *Hermetia illucens*

**DOI:** 10.1101/2023.06.20.544772

**Authors:** Frédéric Manas, Carole Labrousse, Christophe Bressac

## Abstract

In polyandrous species, competition between males for offspring paternity goes on after copulation through the competition of their ejaculates for the fertilisation of female’s oocytes. Given that males allocating more spermatozoa are favored, different models of sperm competition predict adaptive plasticity in male sperm production and allocation. These predictions were tested experimentally in the black soldier fly (BSF) *Hermetia illucens*. In this farmed insect, adult biology is little known despite the economic interest of larvae for bioconversion and as an animal feedstuff. Two sets of experiments were carried out to modify the risk of sperm competition perceived by males. The first consisted of placing adult males alone or in groups of 10 – modifying mean risk of sperm competition – and then measuring their sperm production. The second took place at the beginning of copulation; pairs with males from the two mean risk of sperm competition treatments were transferred to different contexts of immediate risk of sperm competition (empty cages, cages containing 10 males, or cages containing 10 females) and the number of spermatozoa stored by the females was counted. Males reared in groups of 10 showed more spermatozoa in their seminal vesicles than males reared alone. Regarding sperm allocation, females that mated in the presence of conspecifics – either 10 males or 10 females – stored more spermatozoa than those that mated alone. This study shows that sperm production and allocation are dependent on sperm competition risk in BSF, revealing a plasticity of reproduction under socio-sexual situations.

## Introduction

The struggle for reproduction is an important selective pressure leading to many evolutionary adaptations which is particularly typified by the competition between males of polyandrous species (Andersson, 1994). Fifty years ago, Parker (1970) theorized that intrasexual competition between males could be expressed both before and after copulation, as it could continue within female reproductive organs, in the form of sperm competition - i.e. ‘the competition within a single female between the sperm from two or more males over the fertilization of the ova’.

Many physiological (Pizzari & Parker, 2009; Godwin et al., 2017), morphological (Córdoba-Aguilar et al., 2003) and behavioral (Alcock, 1994; Cueva del Castillo, 2003; Barbosa, 2012) traits have been interpreted in light of this paradigm shift (Parker et al., 1998; Wigby and Chapman, 2004). For example, longer spermatozoa swimming faster, bigger testes producing more spermatozoa or mate-guarding strategies are selected by sperm competition as they maximize male’s fertilization success in the competition (Alcock, 1994; LaMunyon & Samuel, 1999; Godwin et al., 2017; Lüpold et al., 2020). Among these traits, plasticity in sperm production (i.e. spermatogenesis) and sperm allocated to each copulation have been the subject of many predictions (Parker, 1970; Parker et al., 1997; Parker & Pizzari, 2010). Based on the costs to males of spermatozoa and seminal fluid contents (Dewsbury, 1982; Olsson et al., 1997), theoretical models predict fitness benefits when males strategically adjust ejaculate expenditure depending on the mating context (Parker & Pizzari, 2010). These models are basically divided into two categories : those based on the “risk” of sperm competition (Parker, 1990; Parker et al., 1997) and those based on the “intensity” of sperm competition (Ball & Parker, 1996, 1997). The risk of sperm competition is defined as the probability that the sperm of a male will compete with the sperm of other males for fertilization of a defined set of ova (Parker, 1990; Parker et al., 1997) whereas the intensity of sperm competition, usually applied to external fertiliser, is the actual number of ejaculates competing for a given set of eggs (Ball & Parker, 1996, 1997).

The predictions of “risk” models have been successfully tested in many organisms, including rodents, fish, and many insects (delBarco-Trillo, 2011). For example, in *Drosophila melanogaster*, sperm production increases when males are housed with other males for a long period of time (Moatt et al., 2014) – increasing what has been named “mean risk of sperm competition” (Engqvist & Reinhold, 2005). Moreover, the arrival of rival males during copulation – increasing the “immediate risk of sperm competition” (Engqvist & Reinhold, 2005) - induces focal males to transfer more spermatozoa to the female (Garbaczewska et al., 2013). On the contrary, “intensity” models predictions are more challenging to test and little evidence is available as to the veracity of their predictions (Kelly & Jennions, 2011 but see Sloan et al., 2018).

The quantity of sperm produced or allocated is not the only component of copulation modified according to the context of sperm competition. For instance, the duration of copulation is particularly studied as it can be considered as a proxy for the amount of sperm allocated (Bretman et al., 2009; Barbosa, 2011), although it is not always true (see Weggelaar et al., 2019). Regardless of the sperm allocation, copulation duration is also predicted to vary with sperm competition risk (Alcock, 1994). By copulating longer, males undertake mate guarding, thus preventing the female from remating (Alcock, 1994), which is a widespread behavior in insects (Lorch et al., 1993; Cueva del Castillo, 2003; Barbosa, 2011).

In this study, we aimed to test the predictions of risk of sperm competition models in the black soldier fly *Hermetia illucens* (BSF), a species of great interest for mass-rearing and organic waste bioconversion (Tomberlin & van Huis, 2020). More precisely, we tested wether males adjust their spermatozoa production and allocation in response to the mean and immediate risk of sperm competition. Despite its economic interest, studies on adults BSF and their reproductive biology are scarce. Giunti et al. (2018) reported a high prevalence of same-sex sexual behaviors in adults BSF, which can be associated to a high degree of polygyny in other species (MacFarlane et al., 2010). Multiple matings have been reported (Permana et al., 2020; Hoffmann et al., 2021) and morphological traits including complex spermathecae, long and numerous spermatozoa, large testes (Munsch-Masset et al., 2023) strongly suggest post-copulatory sexual selection pressures in this species. Here, we experimentally manipulated the risk of sperm competition to examine the phenotypic plasticity in ejaculate expenditure. First, we tested whether long-term exposure to other males could affect sperm production (mean risk of sperm competition) in males’ seminal vesicles. Secondly, we assessed if the sudden appearance or disappearance of rivals (immediate risk of sperm competition) coupled with different mean sperm competition treatments could affect the duration of copulation and the amount of sperm stored in the spermathecae of females.

## Materials and methods

### Rearing conditions

Males and females tested in this study came from a mass rearing maintained in lab conditions since 2018. The strain used is a mix from 2 french commercial lines (Innovafeed and Biomimetic) and one lab line from Wageningen. Black soldier flies were reared under controlled conditions. Adults – ca 200 to 300 - were hosted in 50x50x50 cm cages at 24°C and were provided with a cotton ball saturated with water to maintain moisture. They were exposed to a 12 hours day/night regimen with Philips TLD 36W-84 fluorescent tubes positioned at 10 cm from the cages and providing 2000 to 6000 lux. After collection from the rearing cage, eggs and larvae were maintained at 27°C, the developing substrate was the Gainesville diet (Tomberlin & Sheppard, 2002), no additional moisture was added during development. Pupae were collected and maintained in groups at 24°C with sawdust until emergence. Emerging flies were collected and sexed daily for experiments. Females were isolated in 15x15x15cm cages in groups of 20 females per cage. As for males, they were isolated differently depending on the treatment (see below). Both sexes had access to a cotton ball saturated with water before the experiment. When flies were not used for experiments, they were transferred to the rearing cage.

### Production of spermatozoa

Males of same age (n = 19) were maintained under low risk of sperm competition situation. These individuals were singled, placed in individual 120 mL plastic containers preventing any visual or physical contact with other males and limiting olfactory cues. The second treatment consisted in placing ten males of the same age in a 960 mL plastic container allowing physical, visual, and chemosensory contacts, to simulate a high risk of sperm competition (n = 24 individuals dissected). Males from both treatments were kept in these conditions from 5 to 8 days with access to a cotton ball saturated with water.

### Allocation of spermatozoa

As BSF will not initiate copulations when only one male and one female are placed in a cage (personal observations), the first step of the experiment on the immediate risk of sperm competition involved transferring 20 virgin males from both treatments (10 singled males and 10 grouped males) to one 15x15x15 cm cage containing 20 virgin females. Males from the singled treatment and from the grouped treatment were distinguished by the application of distinct colored spots on their thorax.

Individuals remained in contact for 5 hours and fourteen observations sessions were performed. The cage was constantly watched during the contact time to ensure that only first matings for each individual were recorded.

Once copulations began, each mating pair was gently placed on the lid of a petri dish and transferred in a cage of similar size containing either no individuals to simulate low immediate risk of sperm competition (n = 38 individuals composed by n = 23 grouped males and n = 15 singled males), 10 males to simulate high immediate risk of sperm competition (n = 38 individuals composed by n = 24 grouped males and n = 14 singled males), or 10 females to test the ability of males to recognise genuine competitors (n = 43 individuals composed by n = 27 grouped males and n = 16 singled males). Time was recorded once copulations were completed, and ‘high risk’ pairs – i.e. pairs that mated in the presence of conspecifics - were removed from the cage immediately after separation to ensure that female did not re-mate. Pairs were kept together in petri dishes where there is not enough space for extra copulations to occur (see Munsch-Masset et al., 2023), until dissection of the female reproductive tract.

### Dissections and collection of data

Since age can affect the number of spermatozoa in seminal vesicles (Munsch-Masset et al., 2023), we dissected males of similar ages (n = 5 males of 5 days, n = 28 males of 6 days and n = 10 males of 8 days after emergence, and we controlled the age in the statistic models, see below). Dissections were performed under a Nikon SMZ745T stereomicroscope (x3.35 magnification) (Nikon, Japan) in phosphate-buffered saline (PBS) using fine forceps. For all males, the abdomen was opened after decapitation to collect seminal vesicles which were then placed on a slide in PBS and gently uncoiled with fine forceps. Males were photographed under a Nikon SMZ745T stereomicroscope (x3.35 magnification) (Nikon, Japan) with a Leica IC 80 HD camera (Leica, Germany) to measure their head width using ImageJ. This measure can be considered as a reliable proxy of the size of the individuals (Jones & Tomberlin, 2021). In the same way, the seminal vesicles were photographed and their whole length was measured with ImageJ. A drop of DAPI was then applied to the preparation to label the nuclei of the spermatozoa for blind-counting in a section of one of the two seminal vesicles using a fluorescence microscope (Olympus CX40, Japan) with a x20 objective as in Munsch-Masset et al. (2023). The length of this section was also measured to obtain the ratio between the sperm-counted-section and the whole seminal vesicles. Then, this ratio was multiplied to the number of sperm counted within the portion to obtain the total number of spermatozoa in the seminal vesicles. Finally, this was doubled for the total number of sperm of one male.

The dissection of females took place the day after copulation, knowing that females lay eggs two/three days after mating (Munsch-Masset et al., 2023). The two individuals of a pair were photographed under a Nikon SMZ745T stereomicroscope (x3.35 magnification) (Nikon, Japan) with a Leica IC 80 HD camera (Leica, Germany) to measure their head width using ImageJ like the males from the mean risk experiment. These measures were taken into account in the models for both the sperm production and allocation as covariates. For all individuals, the abdomen was opened to collect the three spermathecae which were then placed on a microscope slide. Before crushing them with a microscope coverslip to release the spermatozoa, a drop of DAPI was applied to the spermatheca to mark the nucleus of the spermatozoa which were blind-counted under a fluorescence microscope (Olympus CX40, Japan) under a x20 objective.

### Statistical analyses

To test our hypotheses, linear mixed models (LMM) were used with the “lmer” function in the “lme4” package in R (Bates et al., 2015). We included the head size, the length of the seminal vesicles and the age of the male as covariates and the mean risk of sperm competition treatment as a fixed effect to explain the number of spermatozoa in the seminal vesicles. For the mating experiment, we included the head size of the female, the head size of the male, the age of the pair, copulation duration as covariates and the two risk of sperm competition treatments – mean risk and immediate risk – as well as their interaction as fixed effects to explain the number of spermatozoa in the spermathecae. For the latter, we also added the number of individuals in the cage prior to the begining of mating as it could influence sperm allocation. We applied a logarithmic transformation on the response variables as they were counting data (spermatozoa counts in the seminal vesicles and in the spermathecae). To study the copulation duration, we used cox proportional hazard model with the “coxph” function in the “survival package” in R (Therneau, 2019). In the same way, age, sizes of the male and the female and both sperm competition risk treatments were included as fixed effects in the model.

The day of sampling was included as a random effect to account for variability inherent to each series in the sperm count in the models. The fixed effects in our models were tested using the “lmerTest” package (Kuznetsova et al., 2017), with type III ANOVA F statistics using Satterthwaite approximations for the linear mix models and with type III ANOVA Chi statistics for the survival model. The assumptions of the linear mixed model, including normality of residuals, constant variance, and absence of multicollinearity among the independent variables were checked graphically. We also assessed the proportional hazards assumption of the cox model using Schoenfeld residuals and found no significant violations of this assumption.

All statistical analyses were performed using R version 4.0.2 (R Core Team, 2020). The significance level was set at alpha = 0.05 for all tests. Quantitative data are presented as means ± standard errors (SE) and hazard ratios (HR) are reported for cox models.

## Results

On average, 5 hours of contact at this population density – 20 females and 20 males - allowed 7 ± 0.72 mating to occur, meaning that 35 % actually mated. Through the different observations, grouped males copulated significantly more than singled males (Fisher’s exact test: P = 0.03). In total n = 45 copulations from singled males and n = 74 copulations from grouped males and were observed.

### Production of spermatozoa

The number of spermatozoa found in the seminal vesicles of the males was neither related to their size (F_1,38_ = 0.61 ; P = 0.44, β ± SE = -0.21 ± 0.28) nor with their age (F_1,38_ = 0.62 ; P = 0.44, β ± SE = 0.06 ± 0.08) nor with the length of their seminal vesicles (F_1,38_ = 1.81 ; P = 0.19, β ± SE = 0.07 ± 0.05). However, the treatment of mean risk of sperm competition showed a significant effect on the number of spermatozoa in the seminal vesicles of males (F_1,38_ = 7.96 ; P < 0.01 ; full model R^2^ = 0.20) (Fig.1). Males kept in groups had a mean of 43 % increase in the number of spermatozoa (mean ± SE : 15578 ± 1105, n = 24) in their seminal vesicles compared to singled males (mean ± SE : 10920 ± 1200, n = 19).

**Figure 1-.**
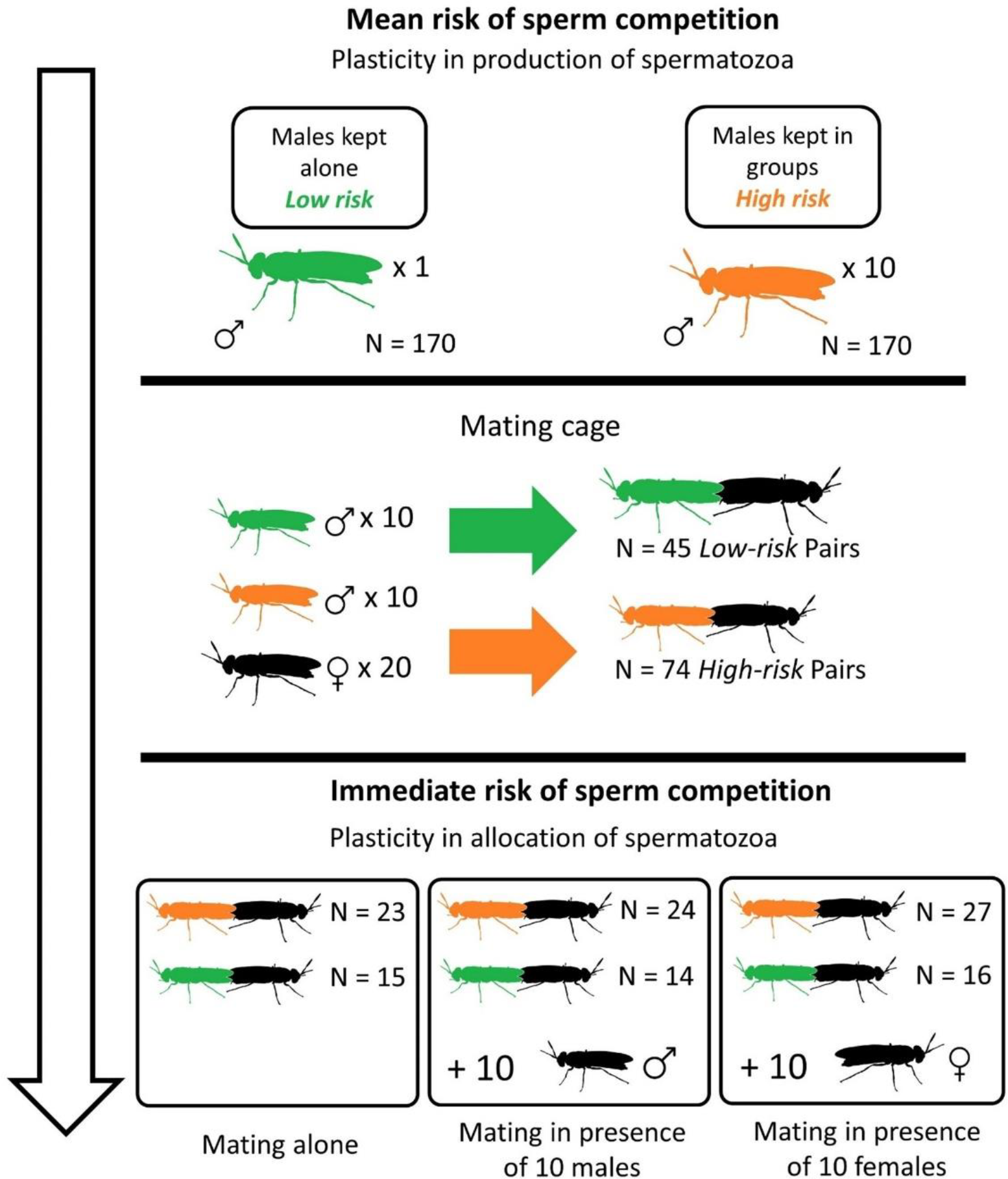
Procedure for the mating experiment to measure the allocation of spermatozoa. Males from two mean risk of sperm competition treatments were transferred to a mating cage. After the begining of mating, the pair was transferred to a different cage according to the immediate risk of sperm competition treatment.

**Figure 2-.**
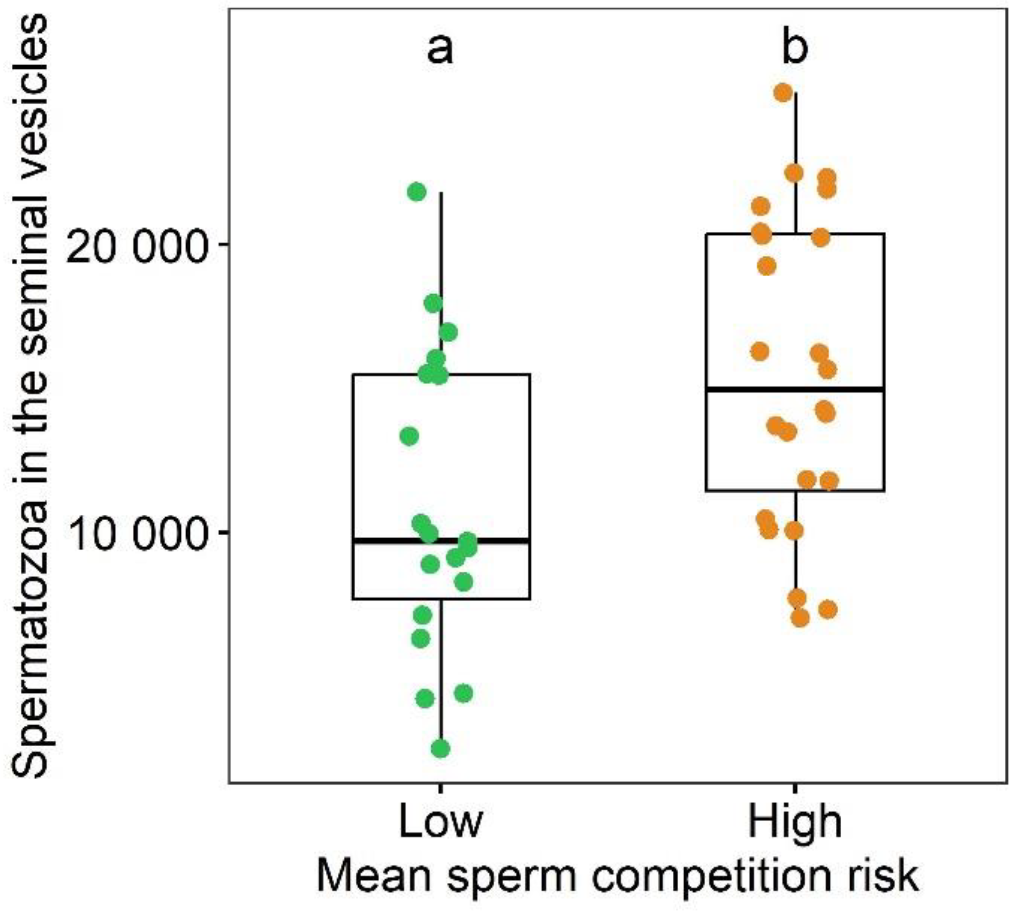
The number of spermatozoa in the seminal vesicles of males according to the mean risk of sperm competition treatment (either the male singled or the male within a group of 10 males) Box plots show median (horizontal bars), upper, and lower quartiles (borders of the box). Whiskers extend from the 10th to the 90th percentiles. Different letters indicate a significant difference at α = 0.05.

### Allocation of spermatozoa

The number of spermatozoa found in the female’s spermathecae was neither related to the size of the male (F_1,107.7_ = 0.61 ; P = 0.44, β ± SE = 0.15 ± 0.19), nor with their age (F_3, 14.1_ = 0.07; P = 0.78; β ± SE = -0.02 ± 0.09), nor to the copulation duration (F_1,104.02_ = 0.01 ; P = 0.96; β ± SE = 0.01 ± 0.01) (Fig.3). The number of spermatozoa in the female’s spermathecae was marginally and positively correlated with the number of individuals in the cage before the beginning of mating (F_1,104.07_ = 3.81 ; P = 0.05; β ± SE = 0.01 ± 0.01). Timing of mating was not different between singled and grouped males (W = 1465, P = 0.27, singled males mean ± SE = 33.6 ± 0.79, grouped males ± SE = 32.32 ± 0.66).

**Figure 3-.**
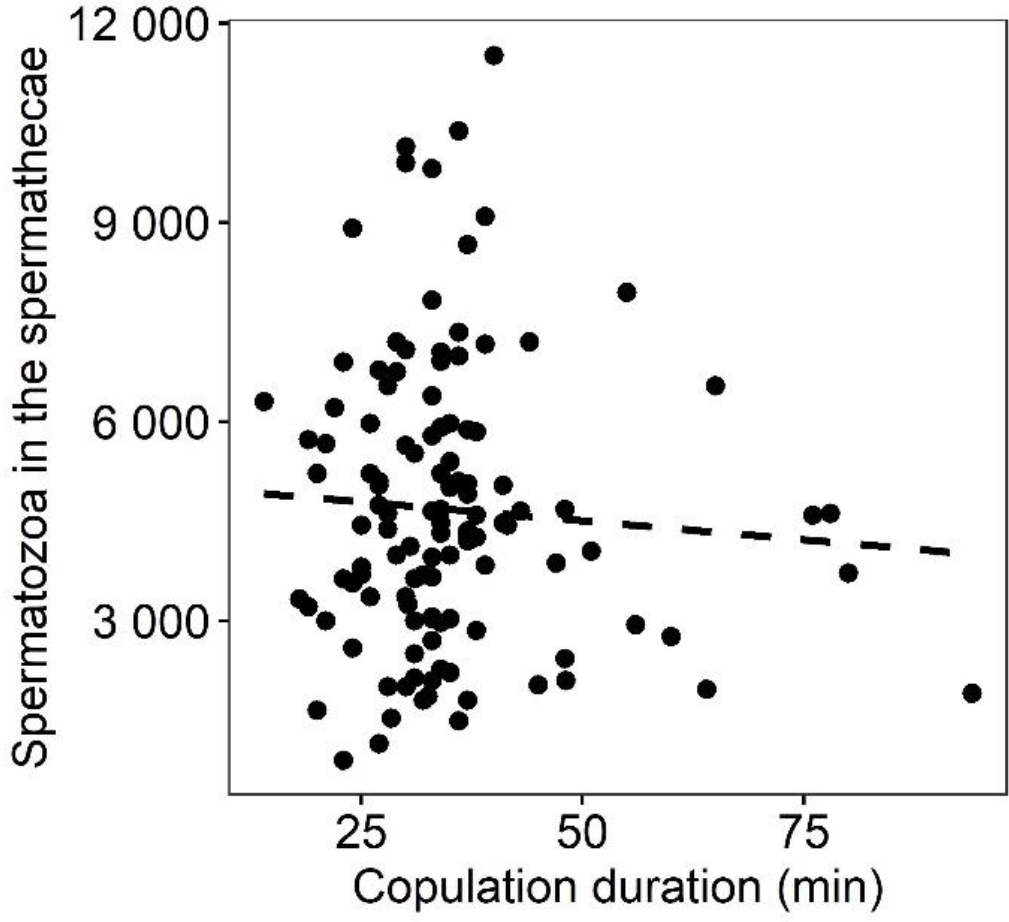
The number of spermatozoa in the spermathecae of females according to the copulation duration. Each point is an individual female, n = 119. The dashed line represents a non significant relationship between these two variables, linear regression: R^2^ < 0.01).

Females that mated with males from the two mean sperm competition risk treatments stored the same amount of spermatozoa (F_1,105.30_= 0.23 ; P = 0.63, Low treatment males : 4614± 273, High treatment males : 4709 ± 270) and the interaction bewteen the mean and the immediate risk treatments was not significant (F_2,103.11_ = 0.08; P = 0.92).

However, the number of spermatozoa found in the female’s spermathecae was related to immediate sperm competition risk treatment (F_2,99.56_ = 6.8; P < 0.01; full model R^2^ = 0.43) (Fig.4). Females that mated in empty cages stored less spermatozoa than the ones that mated in the presence of males (β ± SE = -0.38 ± 0.13; t = -2.84; P < 0.01) and the ones that mated in the presence of females (β ± SE = -0.43 ± 0.13; t = - 3.29; P < 0.01).There was no significative difference (t = -0.37, P = 0.71) between the content of spermathecae of females mated with males in the 10 males treatment (mean ± SE : 4943 ± 376, n = 38) and those in the 10 females treatment (mean ± SE : 5554 ± 284, n = 43). Females mated with males in the presence of conspecifics – either males or females - had a mean 60 % increase in the number of spermatozoa (mean ± SE : 5268 ± 233, n = 81 with n = 38 with pairs 10 males and n = 43 pairs with 10 females) compared to the pairs mating alone (mean ± SE : 3406 ± 268, n = 38).

**Figure 4-.**
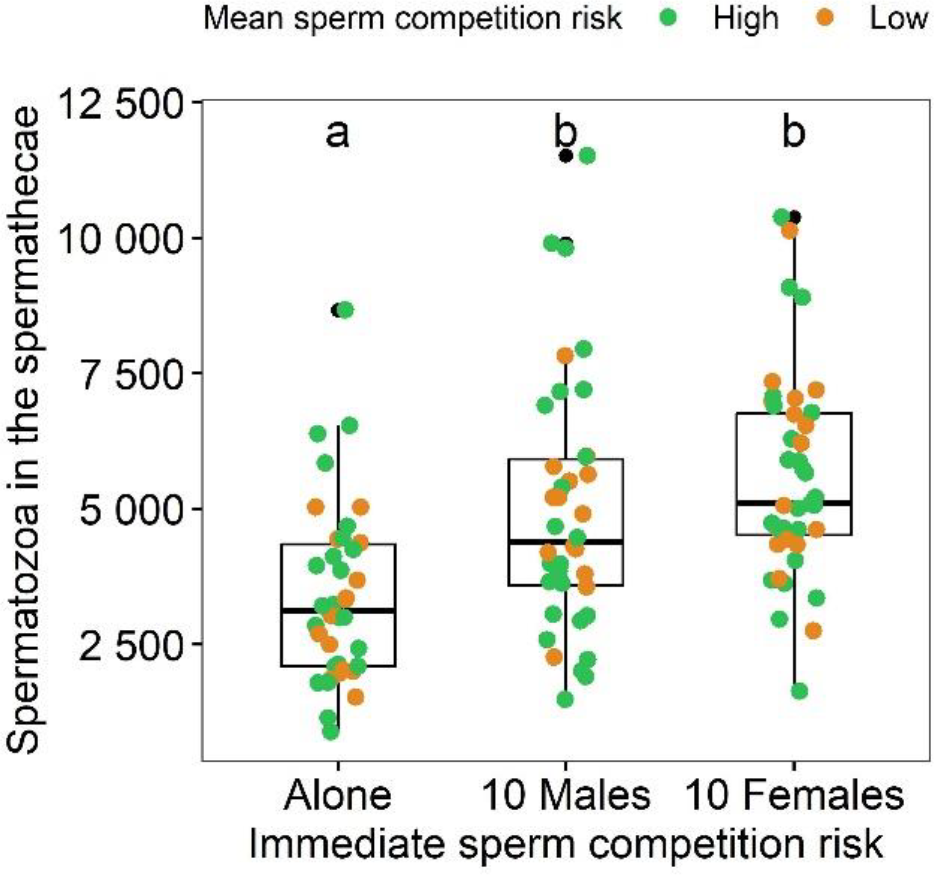
The number of spermatozoa in the spermathecae of females according to the immediate risk of sperm competition (either the pair mating alone, with 10 males or with 10 females). The colors represent the mean sperm competition risk treatment (singled males in green and grouped-males in orange).Box plots show median (horizontal bars), upper, and lower quartiles (borders of the box). Whiskers extend from the 10th to the 90th percentiles. Different letters indicate a significant difference at α = 0.05.

### Copulation duration

The mean time of copulation for all pairs was mean ± SE = 35.16 ± 1.19. Immediate sperm competition risk (χ^2^ = 0.88 ; P = 0.64), mean sperm competition risk treatments (χ^2^ = 0.09 ; P = 0.76) and their interaction (χ^2^ = 3.43 ; P = 0.18) showed no effects on copulation duration. In the same way, the size of the female (χ^2^ = 0.31 ; P = 0.58), the size of the male (χ^2^ = 0.60 ; P = 0.44) and age of the pair (χ^2^ = 1.32 ; P = 0.25) were not related to copulation duration.

## Discussion

BSF males had more spermatozoa in their seminal vesicles when they were grouped, and females of pairs that mated in the presence of conspecifics stored more spermatozoa in their spermathecae. In line with the predictions of the sperm competition theory, males of the BSF respond, on one hand, to mean risk of sperm competition (long-term exposure to rivals) by producing more spermatozoa in their seminal vesicles and on the other hand, to immediate risk of sperm competition (sudden exposure to rivals) by allocating more spermatozoa in a copulation. In contrast, copulation duration was neither related to sperm competition risk treatments, nor to the number of transferred spermatozoa.

Regardless of sperm competition risk, it has been shown that ejaculate expenditure could be condition dependent (Perry & Rowe, 2010; Kaldun & Otti, 2016; Wylde et al., 2020), or sometimes associated with secondary sexual signals (Mautz et al., 2013; Polak et al., 2021). It seems not to be the case in the BSF, in which it has already been demonstrated that male size does not affect sperm production (Jones & Tomberlin, 2021; Munsch-Masset et al., 2023). In the same way, we show here that it does not affect sperm allocation. Interestingly, it appears that males producing more spermatozoa (reared under high risk of sperm competition) do not transfer more spermatozoa to females. Although it is not the case, one would expect that the amount of spermatozoa available to males might be partly determinant of the amount allocated to a copulation (Engqvist & Reinhold, 2005) and finally stored by females. For the latter, it is impossible to disentangle the role of the male from the role of the female on the storage of spermatozoa. We cannot exclude the male mating rate hypothesis (Vahed & Parker, 2012) - i.e. the possibility that the increase in sperm production in group-reared males is due to a higher estimate of the number of mating opportunities based on density. Such an increase in sperm production could be explained by the risk of sperm competition or by the number of opportunities for successive mating, although these two explanations are not mutually exclusive. Furthermore, while allocating the same quantity of spermatozoa to a unique mating (see results), we observed that grouped males had higher success in mating than singled males, suggesting that density induce a greater number of copulation attempts. To investigate this hypothesis further, a different experiment would have to be set up to allow the possibility of multiple copulations per treatment.

In contrast to some other species like the yellow dung fly *Scatophaga stercoraria* (Simmons et al., 1999) or the scorpionfly *Panorpa cognata* (Engqvist & Sauer, 2003), copulation duration is not related to the amount of sperm transferred by the male in BSF. Sperm transfer dynamics that do not follow a linear relationship with time are not rare (Weggelaar et al., 2019) and it seems to be the case in the black soldier fly. Here, durations of copulations were not different between the three treatments. Apart from sperm transfer dynamics, plasticity in copulation duration when males are exposed to rivals can be associated with active mate-guarding (Lorch et al., 1993; Alcock, 1994; Cueva del Castillo, 2003), a behavior that BSF males do not appear to exhibit (Giunti et al., 2018), as confirmed here. Also, it would appear that once copulation has begun, surrounding males lose interest in the pair, unlike during courtship when challenger males may pounce on the pair attempting copulation (Julita et al., 2020).

Numerous cues can be used by males to assess the risk of sperm competition. For example, another Diptera, *Drosophila melanogaster* uses combinations of cues as diverse as visual, contacts, chemosensory, and sounds to detect rivals (Bretman et al., 2011). It has been suggested that BSF uses acoustic signals to identify conspecifics without differentiating females from potential rivals (Giunti et al., 2018), leading frequent same-sex sexual behaviors. These behaviors are observed with males displaying aedeagus eversion during courtship (Personal observations, Giunti et al., 2018), which may indicate that males of the BSF attempt to copulate indifferently with males and females. Same-sex sexual behaviors are common in species with high density of individuals where females are hard to distinguish (Scharf & Martin, 2013) as in BSF rearings, but wether these are current in the wild is not known. Interestingly, we found that BSF males appeared to adjust the number of spermatozoa allocated in a copulation when they were with conspecifics, regardless of whether these were males or females. Although this effect may be due to an effect of population density, this sperm adjustment is in line with a potential absence of sex recognition in BSF males.

Like many aspects of BSF biology, pre-copulatory sexual selection processes in this species are not precisely known. Sexual dimorphism is reduced and preliminary results indicate that male size does not play a role in female’s mates selection (our unpublished data). BSF was described as using leks to mate (Tomberlin & Sheppard, 2001). Those structures are defined as aggregated males display sites that females attend primarily for the purpose of fertilization (Höglund & Alatalo, 1995). Supposedly aggressive intrasexual interactions were also observed but females were said to be ‘similarly greeted’ than males in the supposed lek sites, except that these interactions ended in matings (Tomberlin & Sheppard, 2001). We did not notice any aggregating area akin to a lek in our rearing conditions (Benelli et al., 2014), furthermore the possible lack of sex recognition may question the hypothesis of the BSF actually being a lekking species. Previous studies have demonstrated the occurrence of multiple mating in BSF (Permana et al., 2020; Hoffmann et al., 2021). Consistently with sperm competition theory, our findings suggest that males invest more in sperm production and allocation as a strategy to overcome rivals in this competitive reproductive environment. However, a bet hedging strategy is not evidenced here because males copulating in the presence of virgin females do not spare their sperm reserves in the perspective of the insemination of a maximum number of mates. Besides sperm competition, the complexity of female spermathecae in this species (Munsch-Masset et al., 2023) strongly suggests that post-copulatory intersexual selection mechanisms are at work, such as cryptic female choice or sperm precedence (Pascini & Martins, 2017).

BSF is a species of great economic interest in animal production for its potential as a feed source (Tomberlin & van Huis, 2020). Rearing conditions are certainly very different from natural conditions which are still little known. Not all individuals in a breeding mate, and only a proportion of them mate several times (our unpublished data). When it comes to controlling reproduction, for example to make genetic selection, understanding sexual selection processes is a key factor that needs to be integrated in the rearing methods – e.g., by determining whether different contexts of competition are good for productivity.

## Acknowledgments

We thank Elisabeth Herniou and Harmony Piterois for the proofreading of the paper. We thank Hélène Girotvergne for technical assistance. This revised version was greatly improved by suggestions of Rebecca Boulton, Isabel Smallegange and one anonymous reviewer.

## Funding

FM was funded by the Doctoral School ‘Santé, Sciences Biologiques et Chimie du Vivant’. This work is part of the BioSexFly program funded by the Centre Val de Loire region.

## Conflict of interest disclosure

The authors declare that they comply with the PCI rule of having no financial conflicts of interest in relation to the content of the article.

## Data, scripts, code, and supplementary information availability

Analyses reported in this article can be reproduced using the data and script provided by Frédéric Manas (2023) (https://zenodo.org/records/10078561). https://doi.org/10.5281/zenodo.10488164

## Notes

https://doi.org/10.5281/zenodo.10488164

### Competing Interest Statement

The authors have declared no competing interest.

### Summary of Updates

We changed the title of the article in biorxiv

## References

Alcock J (1994) Postinsemination Associations Between Males and Females in Insects: The Mate-Guarding Hypothesis. Annual Review of Entomology, 39, 1–21. 10.1146/annurev.en.39.010194.000245

Andersson M (1994) Sexual selection. Princeton University Press.

Ball MA, Parker GA (1996) Sperm Competition Games: External Fertilization and “Adapative” Infertility. Journal of Theoretical Biology, 180, 141–150. 10.1006/jtbi.1996.0090

Ball MA, Parker GA (1997) Sperm Competition Games: Inter- and Intra-species Results of a Continuous External Fertilization Model. Journal of Theoretical Biology, 186, 459–466. 10.1006/jtbi.1997.0406

Barbosa F (2011) Copulation duration in the soldier fly: the roles of cryptic male choice and sperm competition risk. Behavioral Ecology, 22, 1332–1336. 10.1093/beheco/arr137

Barbosa F (2012) Males responding to sperm competition cues have higher fertilization success in a soldier fly. Behavioral Ecology, 23, 815–819. 10.1093/beheco/ars035

Bates D, Mächler M, Bolker B, Walker S (2015) Fitting Linear Mixed-Effects Models using lme4. Journal of Statistical Software, 67, 1–48. 10.18637/jss.v067.i01

Benelli G, Daane KM, Canale A, Niu CY, Messing RH, Vargas RI (2014) Sexual communication and related behaviours in Tephritidae: current knowledge and potential applications for Integrated Pest Management. Journal of Pest Science, 87, 385–405. 10.1007/s10340-014-0577-3

Bretman A, Fricke C, Chapman T (2009) Plastic responses of male Drosophila melanogaster to the level of sperm competition increase male reproductive fitness. Proceedings of the Royal Society B: Biological Sciences, 276, 1705–1711. 10.1098/rspb.2008.1878

Bretman A, Westmancoat JD, Gage MJG, Chapman T (2011) Males use multiple, redundant cues to detect mating rivals. Current Biology, 21, 617–622. 10.1016/j.cub.2011.03.008

Córdoba-Aguilar A, Uhía E, Rivera AC (2003) Sperm competition in Odonata (Insecta): the evolution of female sperm storage and rivals’ sperm displacement. Journal of Zoology, 261, 381–398. 10.1017/S0952836903004357

Cueva del Castillo R (2003) Body size and multiple copulations in a neotropical grasshopper with an extraordinary mate-guarding duration. Journal of Insect Behavior, 16, 503–522. 10.1023/A:1027303323242

Dewsbury DA (1982) Ejaculate cost and male choice. The American Naturalist, 119, 601–610. 10.1086/283938

Engqvist L, Reinhold K (2005) Pitfalls in experiments testing predictions from sperm competition theory. Journal of Evolutionary Biology, 18, 116–123. 10.1111/j.1420-9101.2004.00792.x

Engqvist L, Sauer KP (2003) Determinants of sperm transfer in the scorpionfly Panorpa cognata: male variation, female condition and copulation duration. Journal of Evolutionary Biology, 16, 1196–1204. 10.1046/j.1420-9101.2003.00613.x

Garbaczewska M, Billeter JC, Levine JD (2013) Drosophila melanogaster males increase the number of sperm in their ejaculate when perceiving rival males. Journal of Insect Physiology, 59, 306–310. 10.1016/j.jinsphys.2012.08.016

Giunti G, Campolo O, Laudani F, Palmeri V (2018) Male courtship behaviour and potential for female mate choice in the black soldier fly Hermetia illucens L. (Diptera: Stratiomyidae). Entomologia Generalis, 38, 29–46. 10.1127/entomologia/2018/0657

Godwin JL, Vasudeva R, Michalczyk Ł, Martin OY, Lumley AJ, Chapman T, Gage MJG (2017) Experimental evolution reveals that sperm competition intensity selects for longer, more costly sperm. Evolution Letters, 1, 102–113. 10.1002/evl3.13

Hoffmann L, Hull KL, Bierman A, Badenhorst R, Bester-van der Merwe AE, Rhode C (2021) Patterns of genetic diversity and mating systems in a mass-reared black soldier fly colony. Insects, 12, 480. 10.3390/insects12060480

Höglund J, Alatalo RV (1995) Leks: Princeton University Press.

Jones BM, Tomberlin JK (2021) Effects of adult body size on mating success of the black soldier fly, Hermetia illucens (L.) (Diptera: Stratiomyidae). Journal of Insects as Food and Feed, 7, 5–20. 10.3920/JIFF2020.0001

Julita U, Lusianti Fitri L, Eka Putra R, Dana Permana A (2020) Mating success and reproductive behavior of black soldier fly Hermetia illucens l. (Diptera, Stratiomyidae) in tropics. Journal of Entomology, 17, 117–127. 10.3923/je.2020.117.127

Kaldun B, Otti O (2016) Condition-dependent ejaculate production affects male mating behavior in the common bedbug Cimex lectularius. Ecology and Evolution, 6, 2548–2558. 10.1002/ece3.2073

Kelly CD, Jennions MD (2011) Sexual selection and sperm quantity: meta-analyses of strategic ejaculation. Biological Reviews, 86, 863–884. 10.1111/j.1469-185X.2011.00175.x

Kuznetsova A, Brockhoff PB, Christensen RHB (2017) lmerTest Package: Tests in Linear Mixed Effects Models. Journal of Statistical Software, 82. 10.18637/jss.v082.i13

LaMunyon CW, Samuel W (1999) Evolution of sperm size in nematodes: sperm competition favours larger sperm. Proceedings of the Royal Society of London. Series B: Biological Sciences, 266, 263–267. 10.1098/rspb.1999.0631

Lorch Patrick D, Wilkinson Gerald S, Reillo Paul R (1993) Copulation duration and sperm precedence in the stalk-eyed fly Cyrtodiopsis whitei (Diptera : Diopsidae). Behavioral Ecology and Sociobiology, 32. 10.1007/BF00183785

Lüpold S, de Boer RA, Evans JP, Tomkins JL, Fitzpatrick JL (2020) How sperm competition shapes the evolution of testes and sperm: a meta-analysis. Philosophical Transactions of the Royal Society B: Biological Sciences, 375, 20200064. 10.1098/rstb.2020.0064

MacFarlane GR, Blomberg SP, Vasey PL (2010) Homosexual behaviour in birds: frequency of expression is related to parental care disparity between the sexes. Animal Behaviour, 80, 375–390. 10.1016/j.anbehav.2010.05.009

Mautz BS, Møller AP, Jennions MD (2013) Do male secondary sexual characters signal ejaculate quality? A meta-analysis. Biological Reviews, 88, 669–682. 10.1111/brv.12022

Moatt JP, Dytham C, Thom MD (2014) Sperm production responds to perceived sperm competition risk in male Drosophila melanogaster. Physiology & Behavior, 131, 111–114. 10.1016/j.physbeh.2014.04.027

Munsch-Masset P, Labrousse C, Beaugeard L, Bressac C (2023) The reproductive tract of the black soldier fly (Hermetia illucens) is highly differentiated and suggests adaptations to sexual selection. Entomologia Experimentalis et Applicata. 10.1111/eea.13358

Olsson M, Madsen T, Shine R (1997) Is sperm really so cheap? Costs of reproduction in male adders, Vipera berus. Proceedings of the Royal Society of London. Series B: Biological Sciences, 264, 455–459. 10.1098/rspb.1997.0065

Parker GA (1970) Sperm competition and its evolutionary consequences in the insects. Biological Reviews, 45, 525–567. 10.1111/j.1469-185X.1970.tb01176.x

Parker GA (1990) Sperm Competition Games: Raffles and Roles. Proceedings: Biological Sciences, 242, 120–126. 10.1098/rspb.1990.0114

Parker GA, Ball MA, Stockley P, Gage MJG (1997) Sperm competition games: a prospective analysis of risk assessment. Proceedings of the Royal Society of London. Series B: Biological Sciences, 264, 1793–1802. 10.1098/rspb.1997.0249

Parker GA, Birkhead TR, Møller AP (1998) Sperm competition and sexual selection. Sperm competition and the evolution of ejaculates: towards a theory base, 3–54.

Parker GA, Pizzari T (2010) Sperm competition and ejaculate economics. Biological Reviews, 85, 897–934. 10.1111/j.1469-185X.2010.00140.x

Pascini TV, Martins GF (2017) The insect spermatheca: an overview. Zoology, 121, 56–71. 10.1016/j.zool.2016.12.001

Permana AD, Fitri LL, Julita U (2020) Influence of mates virginity on black soldier fly, Hermetia illucens L. (Diptera: stratiomyidae) mating performance. Jurnal Biodjati, 5, 174–181. 10.15575/biodjati.v5i2.9049

Perry JC, Rowe L (2010) Condition-dependent ejaculate size and composition in a ladybird beetle. Proceedings of the Royal Society B: Biological Sciences, 277, 3639–3647. 10.1098/rspb.2010.0810

Pizzari T, Parker GA (2009) Sperm competition and sperm phenotype. In: Sperm Biology, pp. 207–245. lsevier. 10.1016/B978-0-12-372568-4.00006-9

Polak M, Hurtado-Gonzales JL, Benoit JB, Hooker KJ, Tyler F (2021) Positive genetic covariance between male sexual ornamentation and fertilizing capacity. Current Biology, 31, 1547–1554.e5. 10.1016/j.cub.2021.01.046

Scharf I, Martin OY (2013) Same-sex sexual behavior in insects and arachnids: prevalence, causes, and consequences. Behavioral Ecology and Sociobiology, 67, 1719–1730. 10.1007/s00265-013-1610-x

Simmons LW, Parker GA, Stockley P (1999) Sperm Displacement in the Yellow Dung Fly, Scatophaga stercoraria: An Investigation of Male and Female Processes. The American Naturalist, 153, 302–314. 10.1086/303171

Sloan NS, Lovegrove M, Simmons LW (2018) Social manipulation of sperm competition intensity reduces seminal fluid gene expression. Biology Letters, 14, 20170659. 10.1098/rsbl.2017.0659

Therneau T (2019) The survival package. https://github.com/therneau/survival

Tomberlin JK, Van Huis A (2020) Black soldier fly from pest to ‘crown jewel’ of the insects as feed industry: an historical perspective. Journal of Insects as Food and Feed, 6, 1–4. 10.3920/JIFF2020.0003

Tomberlin JK, Sheppard DC (2001) Lekking behavior of the black soldier fly (Diptera: Stratiomyidae). The Florida Entomologist, 84, 729. 10.2307/3496413

Tomberlin JK, Sheppard DC (2002) Factors influencing mating and oviposition of black soldier flies (Diptera: Stratiomyidae) in a colony. Journal of Entomological Science, 37, 345–352. 10.18474/0749-8004-37.4.345

delBarco-Trillo J (2011) Adjustment of sperm allocation under high risk of sperm competition across taxa: a meta-analysis. Journal of Evolutionary Biology, 24, 1706–1714. 10.1111/j.1420-9101.2011.02293.x

Vahed K, Parker DJ (2012) The evolution of large testes: sperm competition or male mating rate? Ethology, 118, 107–117. 10.1111/j.1439-0310.2011.01991.x

Weggelaar TA, Commandeur D, Koene JM (2019) Increased copulation duration does not necessarily reflect a proportional increase in the number of transferred spermatozoa. Animal Biology, 69, 95–115. 10.1163/15707563-00001078

Wigby S, Chapman T (2004) Sperm competition. Current Biology, 14, R100–R103. 10.1016/j.ympev.2004.01.013

Wylde Z, Crean A, Bonduriansky R (2020) Effects of condition and sperm competition risk on sperm allocation and storage in neriid flies. Behavioral Ecology, 31, 202–212. 10.1093/beheco/arz178

